# Private haplotype barcoding facilitates inexpensive high-resolution genotyping of multiparent crosses

**DOI:** 10.1101/116582

**Authors:** Daniel A. Skelly, John H. McCusker, Eric A. Stone, Paul M. Magwene

## Abstract

Inexpensive, high-throughput sequencing has led to the generation of large numbers of sequenced genomes representing diverse lineages in both model and non-model organisms. Such resources are well suited for the creation of new multiparent populations to identify quantitative trait loci that contribute to variation in phenotypes of interest. However, despite significant drops in per-base sequencing costs, the costs of sample handling and library preparation remain high, particularly when many samples are sequenced. We describe a novel method for pooled genotyping of offspring from multiple genetic crosses, such as those that that make up multiparent populations. Our approach, which we call "private haplotype barcoding” (PHB), utilizes private haplotypes to deconvolve patterns of inheritance in individual offspring from mixed pools composed of multiple offspring. We demonstrate the efficacy of this approach by applying the PHB method to whole genome sequencing of 96 segregants from 12 yeast crosses, achieving over a 90% reduction in sample preparation costs relative to non-pooled sequencing. In addition, we implement a hidden Markov model to calculate genotype probabilities for a generic PHB run and a specialized hidden Markov model for the yeast crosses that improves genotyping accuracy by making use of tetrad information. Private haplotype barcoding holds particular promise for facilitating inexpensive genotyping of large pools of offspring in diverse non-model systems.

## Introduction

Advances in high-throughput sequencing have led to remarkable increases in the availability of DNA sequence data from any desired species. This has facilitated the creation of large panels of fully sequenced or densely genotyped individuals from both model and non-model organisms (Keane *et al.* 2011; Cao *et al.* 2011; Wilkening *et al.* 2013; Jeffares *et al.* 2015; Strope *et al.* 2015; The 1000 Genomes Project Consortium 2015). These resources, along with advances in experimental design and the statistical analysis of multiparent populations (e.g.Churchill *et al.* 2004; Scutari *et al.* 2014; Gatti *et al.* 2014;Wei and Xu 2016), present a unique opportunity for identifying quantitative trait loci (QTL) underlying complex traits (Threadgill and Churchill 2012; Huang *et al.* (2015). Indeed, carefully designed multiparent populations are currently being used to understand the genetic architecture of complex trait variation in several major model organisms and crop species (Yu *et al.* 2008; Kover *et al.* 2009;Mackay *et al.* 2012; King *et al.* 2012b; Bandillo *et al.* 2013;Cubillos *et al.* 2013; Mackay *et al.* 2014; Nice *et al.* (2016). These populations show tremendous promise for advancing complex trait genetics, but the cost of their construction is often substantial. Although existing multiparent populations are often justifiably promoted as permanent resources available to all researchers, there is a clear need for methods to facilitate construction of new multiparent populations in non-model organisms with smaller communities or for exploring specific phenotypes in model organisms.

What factors contribute to the costs of constructing and maintaining a multiparent mapping population? Some organism-specific factors such as colony maintenance costs are an unavoidable consequence of biology and can be substantial (Churchill *et al.* 2004). However, the cost of obtaining high-resolution genotypes of offspring represents a large one-time cost required for each multiparent population. In particular, although the costs of high-throughput sequencing have fallen precipitously in recent years (a trend that is likely to continue; https://www.genome.gov/sequencingcosts), costs of sample handling and library preparation have remained comparatively high. When many individuals are sequenced, as for multiparent populations that are well-powered to detect quantitative trait loci (QTL), these library preparation costs represent a significant portion of total genotyping costs (which increase proportionally as sequencing costs decrease).

In this paper, we present a simple method – private haplotype barcoding (PHB) – that utilizes private haplotypes to deconvolve patterns of inheritance in individual offspring from pooled samples. At its core, PHB involves inferring genotypes using short sequencing reads from pooled samples. Short-read sequencing has become a common method for genotyping individuals, often combined with reduced representation approaches which sample a subset of the genome (Baird *et al.* 2008; Andolfatto *et al.* 2011; Elshire *et al.* 2011; Peterson *et al.* 2012). Huang *et al.* (2009)used low coverage whole genome sequencing to genotype recombinant inbred lines from a cross. Andolfatto *et al.* (2011) developed a hidden Markov model (HMM) to infer ancestry in offspring from a backcross genotyped by reduced representation sequencing. King *et al.* (2012a)used an HMM to infer founder ancestry from reduced representation sequencing of members of a multiparent population. Both Xie *et al.* (2010)and Rowan *et al.* (2015) developed HMMs to obtain genotypes of offspring from a cross using low coverage whole genome sequence data. In contrast to all of these approaches, our approach is novel in that we do not barcode samples before pooling and sequencing. This can greatly facilitate sample handling and reduce library preparation costs. Thus, PHB has the potential to drastically lower genotyping costs for certain multiparent population designs. We demonstrate the utility of PHB by obtaining high-resolution genotypes of 96 segregants from 12 yeast crosses, achieving over a 90% reduction in sample preparation costs relative to non-pooled sequencing.

## Results and Discussion

### An overview of strategies for pooled genotyping

A common strategy to lower genotyping costs involves pooling samples and sequencing short reads. Typically, molecular barcodes are employed as a method of distinguishing between distinct sequencing libraries prepared from individual samples but sequenced in a single pool. Known barcode sequences ligated to fragments within each sequencing library allow *in silico* sorting of reads according to their library of origin. Notably, this strategy requires separate library preparation steps for each sample to be genotyped. However, pooling samples for sequencing and *in silico* sorting by sample does not always require molecular barcodes. A major advantage of pooling *without* using barcodes is that samples can be pooled as early as the DNA extraction step, facilitating sample handling and spreading the costs of library preparation across multiple samples. We will refer to this method of pooling as *raw* pooling.

Raw pooling followed by short-read sequencing can be used for many purposes including variant discovery, allele frequency estimation, and sample abundance estimation (Futschik and Schlötterer 2010; Eskin *et al.* 2013). Genotyping individual samples combined in raw pools requires more careful design of the pools, but is still useful in a wide variety of circumstances. For example, barcodes would be unnecessary for genotyping one individual from each of two diverse species with sequenced genomes – say, a human and a fruit fly, where even coding regions show only ~45% sequence identity (Shih *et al.* 2015). In this example, read origin could nearly always be assigned correctly as human or fly because of the high divergence between the species. At another extreme, if one could obtain very long multi-megabase reads with high accuracy, it would be possible to accurately genotype pooled DNA samples of two humans from different geographic regions using known population-specific variation to assign each long read (haplotype) to a sample. Indeed, with infinitely long read lengths and no sequencing errors, only a single marker of unique ancestry on each chromosome would be required to ascertain which chromosomes are found together in a single individual and genotype with perfect fidelity. More generally, the key metric governing success of a raw pooling approach is the fraction of reads overlapping polymorphisms that uniquely distinguish a sample. The space of possible read lengths and minimum pairwise divergence between all samples in a pool can be viewed as a landscape where the feasibility of raw pooling is a joint function of these two parameters (Figure 1).

**Figure 1:**
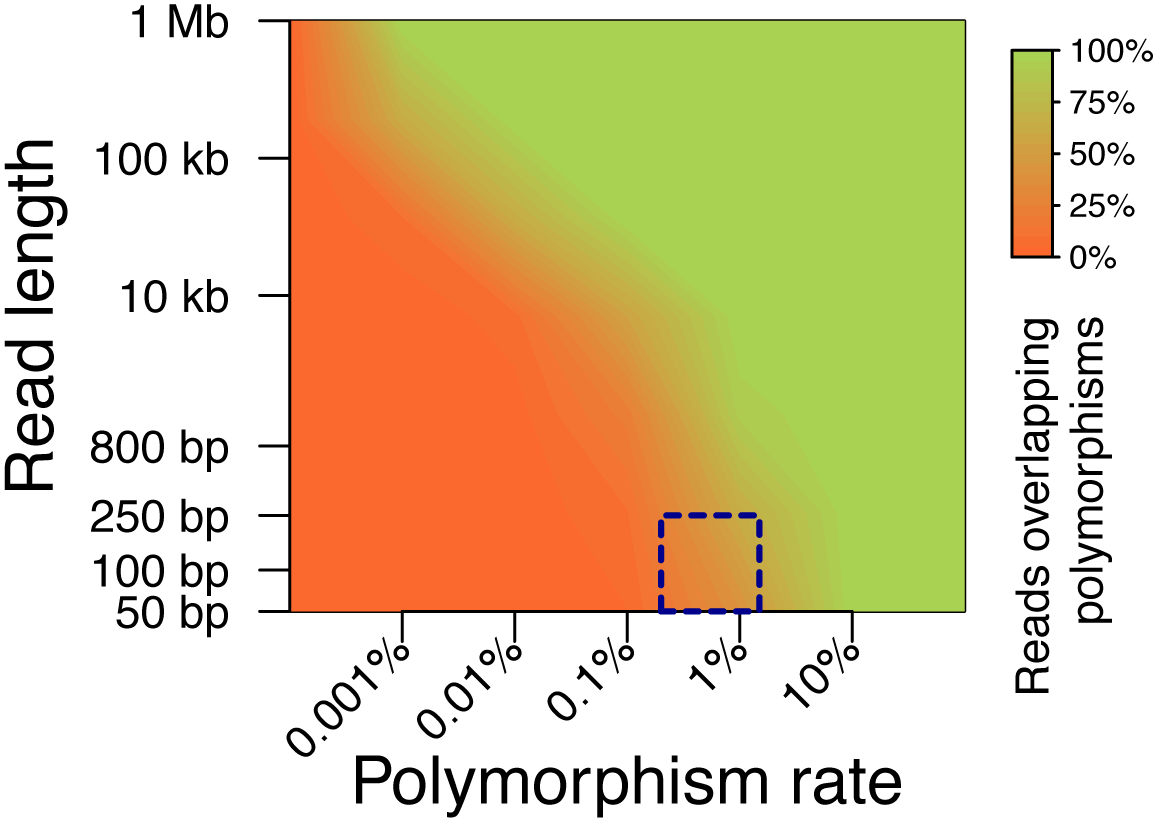
The feasibility of raw pooling as a joint function of read length and pool sequence diversity. Feasibility is quantified as the fraction of reads overlapping polymorphic sites, a significant fraction of which are assumed to be uniquely distinguishing either alone or in combination. For simplicity, we calculate this quantity using a crude model where polymorphisms are modeled as occurring randomly along an infinitely long chromosome. The distance between consecutive polymorphisms is exponentially distributed with rate parameter equal to the polymorphism rate. Then the chance of a read of *N* base pairs overlapping at least one polymorphism is the probability that the distance between any two polymorphisms is less than *N* bases, which is calculated analytically from the exponential distribution function. The box outlined in a dark blue dotted line indicates the approximate region of the plot that is applicable to studies using current short-read technology and populations with levels of diversity that are roughly in line with the yeast strains studied here.

### Private haplotype barcoding reduces sample handling and library preparation costs

The construction of multiparent populations begins with the selection of parental lines, with available high-resolution genotype data, that will contribute ancestry to offspring in the mapping population via a defined breeding scheme. Thus, genotyping offspring that comprise a multiparent mapping population is a simpler task than genotyping a random selection of individuals from a natural population. Specifically, genotyping these offspring can be thought of as tracing the inheritance of segments of the parental genomes in recombinant offspring. PHB is a simple strategy for raw pooling that is particularly useful for genotyping of certain multiparent populations. Because PHB involves genotyping of raw pools, it results in significant reductions in sample handling and library preparation costs in addition to the reduced sequencing costs shared with any pooled genotyping technique.

PHB is a flexible method that may be most clearly illustrated with an example. The strategy involves careful pooling of individuals and computational deconvolution of patterns of haplotype inheritance from the pooled sequencing data. Specifically, we pool samples in such a way that every individual has a portion of ancestry that is unique to the pool (we note that other applications of the method could result in all of each individual’s ancestry being unique to the pool). For example, Figure 2A shows the design of a budding yeast (*Saccharomyces cerevisiae*) multiparent population where each of twelve diverse parental lines are mated to a common “reference” parent and F2 offspring are generated to comprise the mapping population. In this example, one F2 individual from each cross can be combined into a single pool (Figure 2B) and haplotypes inherited from the twelve diverse parental lines can be traced using polymorphisms unique to these parents. Inheritance from the reference parent is inferred to have occurred elsewhere in the genome. This pooling strategy reduces labor and library preparation costs 12-fold in this example.

PHB could also be applied fruitfully for genotyping other multiparent populations or in other quantitative genetic studies. A prerequisite for PHB is that at least part of the genome of each individual in a pool is unique to that pool. Thus, PHB cannot be used to genotype offspring from a multiparent population design where every individual has an equal expected genetic contribution from each founder parent – for example, multiparent recombinant inbred lines such as Collaborative Cross mice or *Arabidopsis thaliana* MAGIC lines (Churchill *et al.* 2004; Kover *et al.* 2009). However, PHB can be used to genotype multiparent populations where individuals that constitute the pool have unique ancestry, and where the non-unique portion of their ancestry can be inferred. Examples of such multiparent populations include the maize nested association mapping population (Yu *et al.* 2008), the *Drosophila* synthetic population resource (King *et al.* 2012a), and the yeast example shown in Figure 2. More broadly, PHB is applicable to breeding designs where individuals are not exchangeable and can be pooled in such a fashion that ancestral haplotypes can be traced to unique individuals in each pool. Although PHB is not limited to applications involving multiparent mapping populations, it may be particularly pertinent for such populations because they tend to be composed of individuals with high genomic sequence diversity and haplotype blocks that are much longer than short read sequence lengths.

**Figure 2:**
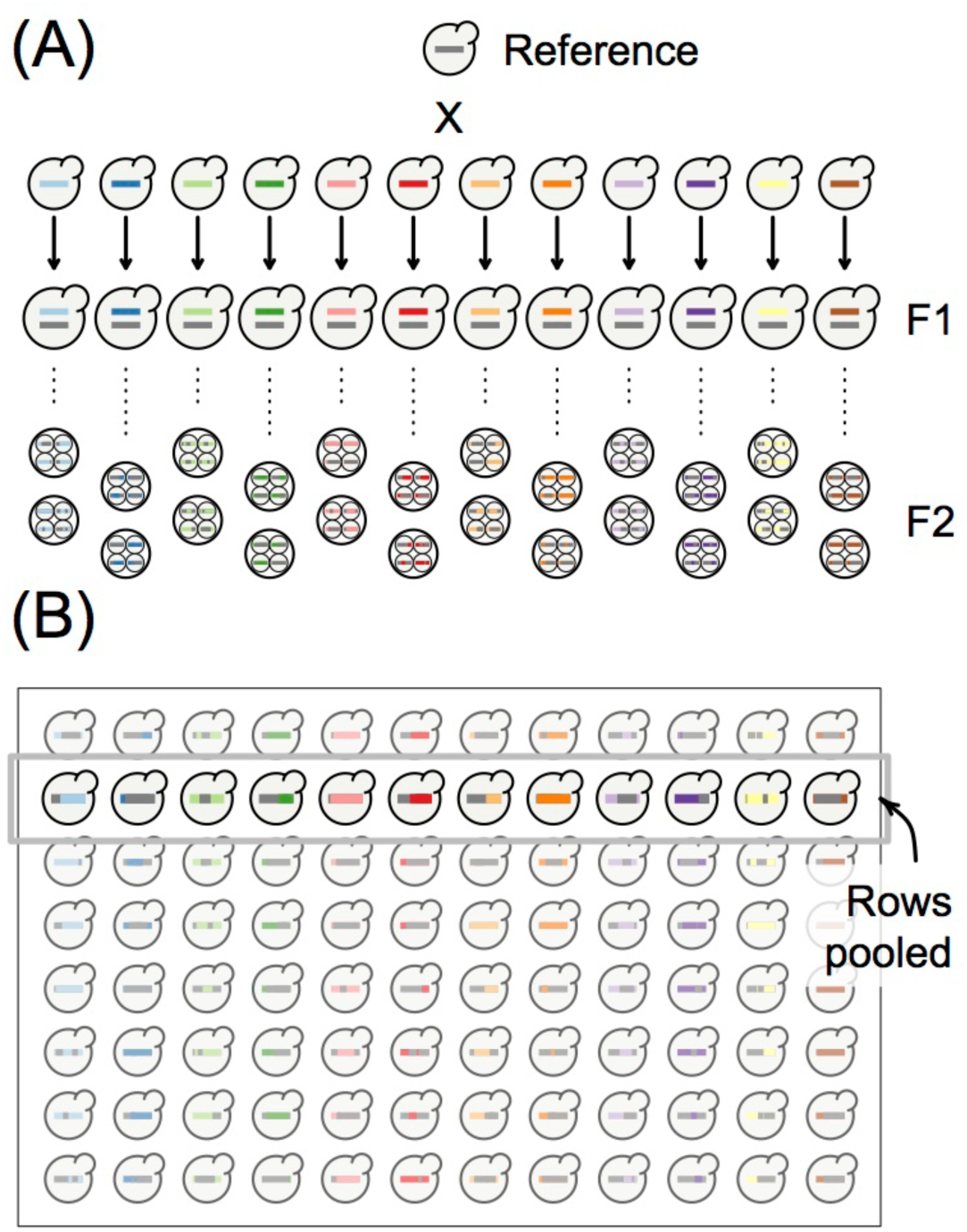
Design of the PHB genotyping study. (A) The breeding design of a yeast multiparent mapping population where each of twelve diverse parental lines are mated to a common “reference” parent and eight F2 offspring are generated from each cross. Large X represents a cross, with arrows leading to a depiction of the resulting diploid hybrid. Dotted line represents meiosis. Offspring are shown as a tetrad, with the four haploid spores that result from a single round of meiosis encapsulated in a sac known as an ascus. Two ascii per cross were analyzed in this study. (B) The design shown in (A) is conveniently suited to plate-based sample handling approaches. To carry out PHB for genotyping, we pooled rows of the plate.

### PHB accurately genotypes yeast segregants

To demonstrate the power of PHB for accurately inferring genotypes from a pooled sequencing library, we obtained 96 yeast segregants according to the breeding design shown in Figure 2A. Specifically, we crossed each of twelve diverse parental lines to a common “reference” parent and generated eight F2 offspring from each cross to comprise the population. This design is conveniently suited to plate-based sample handling approaches (Figure 2B). To carry out PHB to obtain genotypes, we pooled rows of the plate (Figure 2B). Note that, for a given row, each haploid segregant is expected to inherit half of its genome from the common reference parent and the other half from one of the diverse set of twelve other parents (with this latter parent unique to the pool).

After pooling cells from each row of the plate, extracting DNA, and carrying out library preparation for short-read sequencing, we obtained millions of short reads from each pool (Table 1). Each parent contributing ancestry to the population has a fully sequenced, *de novo*-assembled genome (Strope *et al.* 2015). We mapped the short reads to a combined genome of all thirteen parents, with a separate contig for each chromosome in each parent (the *pool genome).* Overall, ~97% of reads aligned to at least one genomic locus in the pool genome, with ~5% mapping uniquely to the reference strain and 0.5–2.5% mapping uniquely to one of the other twelve parental strains (Table 1). The higher rate of reads mapping to the reference genome reflects the fact that all individuals in the pool are expected to inherit half of their genomes from the reference strain. These rates of unique mappings are much lower than the percentage of reads (~86%) that uniquely map to the reference genome alone, as opposed to the pool genome. This reflects the fact that many reads do not overlap haplotypes that are private to a single ancestor of the pool. Nevertheless, there remains substantial information about the ancestry of blocks inherited from parental strains for each individual sequenced as part of a pool.

**Table 1.** Mapping statistics from pooled yeast DNA sequencing

| sample |  | Reads<br>(millions) | Percent of reads that map: |  |  |  |  |  |
| --- | --- | --- | --- | --- | --- | --- | --- | --- |
|  |  |  | to ref | uniquely<br>to ref | to pool<br>genome | uniquely to pool genome |  |  |
|  |  |  |  |  |  | total | ref | nonref<br>range* |
| Pooled | A | 47.5 | 95.5 | 86.2 | 97.6 | 18.1 | 5.2 | 0.5–1.7 |
|  | B | 46.6 | 94.9 | 85.4 | 97.3 | 19.0 | 5.2 | 0.6–2.4 |
|  | C | 48.0 | 95.5 | 85.8 | 97.4 | 18.3 | 5.4 | 0.6–1.8 |
|  | D | 51.8 | 95.6 | 86.3 | 97.9 | 18.5 | 5.5 | 0.6–2.4 |
|  | E | 49.8 | 95.4 | 86.1 | 97.5 | 18.7 | 5.4 | 0.5–2.7 |
|  | F | 48.9 | 95.3 | 86.3 | 97.3 | 18.3 | 5.4 | 0.5–1.8 |
|  | G | 45.6 | 95.5 | 86.6 | 97.6 | 19.2 | 5.2 | 0.6–2.4 |
|  | H | 44.3 | 94.7 | 85.5 | 97.3 | 19.1 | 5.3 | 0.5–2.1 |
| control | A10 | 8.4 | 95.8 | 83.7 | 97.6 | 47.9 | 21.3 | 26.6 |
|  | B7 | 10.2 | 95.3 | 86.7 | 97.2 | 51.0 | 20.8 | 30.1 |
|  | G11 | 14.1 | 95.8 | 87.2 | 96.4 | 62.6 | 25.5 | 37.1 |
|  | H3 | 7.0 | 94.9 | 85.8 | 95.7 | 54.3 | 22.7 | 31.6 |

\*For control samples this column lists the percent of reads mapping uniquely to the single non-reference strain that contributed ancestry to the segregant sequenced

Figure 3A shows coverage of uniquely mapping reads in one pooled library across the genome of strain YJM1701, a parental strain contributing ancestry to exactly one segregant in the pool. It is apparent that large blocks of the genome show high read depth while the remaining blocks are very sparsely covered. The blocks with high read depth are composed of many short reads overlapping YJM1701 haplotypes that are unique to the pool, and thus reflect inheritance of that genomic segment from the YJM1701 parent. Conversely, the sparsely covered blocks have few uniquely mapping reads and reflect ancestry of S288c, the reference strain common to all individuals in the pool.

**Figure 3:**
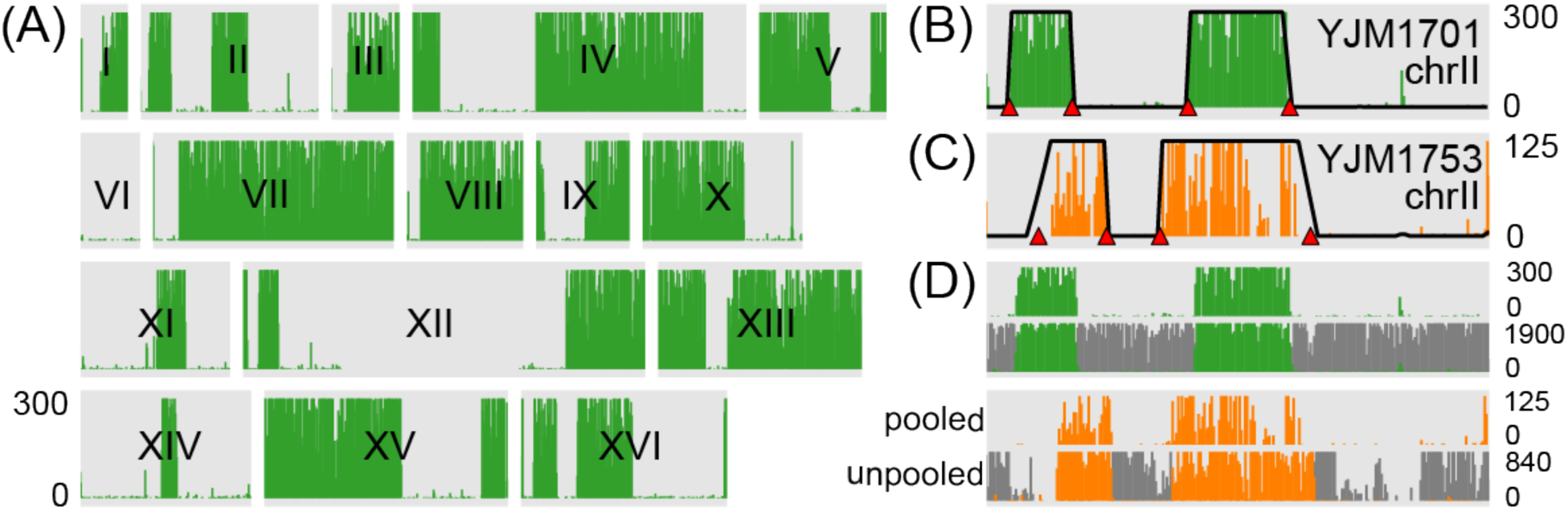
Pooled sequencing read depth reveals blocks of ancestry inherited from reference (YJM1617, isogenic with S288c) or non-reference parental strains. (A) Depth of uniquely mapping reads in 3kb bins tiling across all 16 chromosomes for one spore produced by the YJM1617-YJM1701 hybrid and sequenced in a pooled library. Note the clear boundaries between blocks of ancestry inherited from the non-reference parent. Read depths are trimmed at a maximum of 300 reads as indicated by axis label on the left. Roman numerals indicate individual chromosomes, each bounded by a gray box. The large empty space on chromosome XII is the highly repetitive rDNA array. (B) Detailed view of coverage across chromosome II, with posterior probabilities of nonreference ancestry overlain on each read depth plot. The magnitude of the posterior probability is scaled to the same range as the read depths. Red triangles indicate locations of inferred recombination breakpoints. (C) The same plot as shown in (B), but for a spore with ancestry from a non-reference parent (YJM1753) that is among the least diverged in our pool from the common reference. The read depth threshold is ~40% of that in (A) and (B) but segments of ancestry are still visible. (D) Comparison of ancestry-distinguishing read depth for pooled data versus individually sequenced segregants. Dark grey lines depict ancestry from the reference strain and other colors match subplots (A)-(C). Note much higher read depths in unpooled data.

A naïve approach to inferring ancestry quantitatively would be to simply smooth read coverage across the genome of each non-reference parent unique to the pool and employ a genomic window-based filter to obtain estimates of ancestry originating from reference versus non-reference parents. Instead, we implemented an HMM that provides probabilistic estimates of the genotypes underlying each segregant’s genome. We chose this more sophisticated HMM approach for two reasons. First, the HMM is a way to use the machinery of probability theory to translate from depth of sequencing coverage across a particular bin to a quantitative measure of our confidence in ancestry within that bin. This provides genotype probabilities that would be suitable for QTL mapping. Second, the HMM incorporates information from neighboring bins to inform our confidence in the ancestry calls.

In this HMM, we examine read depth in 7.5kb bins tiling across the genome of each non-reference strain (parent *j)* that contributes ancestry to the pool (i.e. ancestry traceable to one individual in the pool). Following our intuition from above, bins with many reads likely represent a genomic segment inherited from parent *j,* while bins with few reads likely represent either inheritance from the reference parent or bins with low mappability (e.g. due to low sequence complexity). The HMM has two underlying states corresponding to reference ancestry and parent *j* ancestry. Given the total number of reads mapping uniquely to parent *j’s* genome, read depth counts are distributed binomially to each bin. We calibrate bin-specific “success” probabilities using simulation (Methods). In the reference ancestry state, reads are effectively invisible since this ancestry is not unique to the pool, so reads are emitted at bin-specific background error rates. In the parent *j* ancestry state, reads are emitted at higher bin-specific rates. This HMM could be modified to incorporate inference about diploid ancestry in non-inbred diploid individuals, but we do not explore this further as the end product of many multiparent populations is a set of haploid or inbred diploid lines.

This approach allowed us to accurately resolve genotypes of the yeast segregants with high confidence. Figure 3B shows smoothed ancestry probabilities superimposed over read coverage data for chromosome II, demonstrating that estimates of ancestry match intuition. Figure 3C shows data across the same chromosome as Figure 3B but for one of the least diverse non-reference strains in the pool; the similar positions of inherited segments of non-reference ancestry in Figures 3B-C are merely coincidental. Across all yeast segregants that we sequenced, genotype probabilities were strongly bimodal (Figure 4A), with >96% of genomic segments assigned a probability >99% of ancestry from one parent. Thus, genotypes can be confidently assigned as originating from one of two possible haplotypes for the vast majority of the genome.

**Figure 4:**
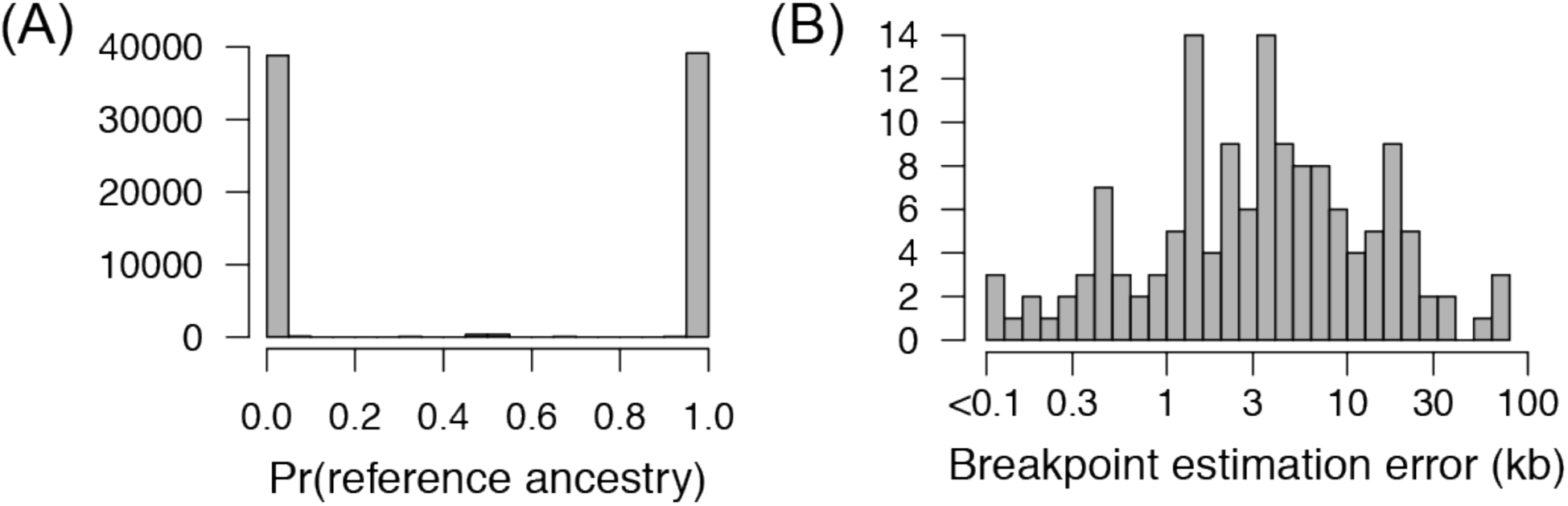
PHB efficiently and accurately infers genotypes of recombinant offspring. (A) Histogram of the probability of reference (YJM1617, isogenic with S288c) ancestry across all 7.5kb bins tiling the yeast genome. This histogram shows probabilities from all pooled segregants sequenced in this study. (B) Histogram of the accuracy of recombination breakpoint estimation. This quantity is measured as the absolute difference between breakpoints inferred using PHB and those calculated by individually sequencing segregants and inferring breakpoint locations. Four segregants were sequenced individually and this histogram combines results across all four.

To objectively assess the accuracy of the genotypes we obtained using PHB, we individually sequenced the genomes *(unpooled* data) of four segregants that were present in PHB pools. We chose the four segregants such that the unique parent of each varied in terms of its “uniqueness” among the thirteen ancestors of the population: one parent was among the most diverse, one was of intermediate diversity, and two were among the least diverse. Figure 3D compares pooled and unpooled data for the segregants shown in Figure 3B-C. Using our unpooled data, we inferred switches in ancestry representing locations of recombination breakpoints, and compared these genomic locations to those inferred using PHB. In general, the locations of breakpoints inferred via PHB were highly accurate, with breakpoints estimated using pooled data within 3.5kb (median) of their estimate in the unpooled data (Figure 4B). Of 166 breakpoints inferred using unpooled data, 141 (87%) were called in our pooled data. Of the 25 missed calls, all fell into one of two categories: (1) two nearby breakpoints constituting a small block of ancestry (median 38kb, *N*=6), or (2) a breakpoint near the end of the chromosome (median 34.6kb from end, *N*=19). There were also 9 breakpoints called in the pooled data not found in the unpooled data, all of which fell into these same two categories. Fortunately, both types of missed calls affect a small portion of the complete genome (for these four segregants, 0.5%–6.1%). Thus, PHB allows us to accurately and efficiently reconstruct genotypes of offspring sequenced in an unlabeled pool.

### Using tetrad structure improves genotyping accuracy and resolution

A peculiarity of yeast biology is that the ascus that contains the four products of a single meiosis (a tetrad) can be physically isolated and each of the four segregants, or spores, can be separated and grown into a colony. Since we tracked spore ancestry when crossing strains to obtain segregants (Figure 2), we developed an alternative HMM for analyzing PHB data that utilizes this tetrad information. Because each tetrad results from a single meiosis, our expectation according to the principles of Mendelian inheritance is that across the tetrad the parental alleles at any genomic locus should show 2:2 segregation in the offspring (Figure 5A, 5B). This HMM jointly models read count data arising from a genomic locus in each of four separate pooled sequencing runs that together contain all the products of twelve separate meioses. The HMM is run separately for each parent (which we will refer to as parent *j*) that uniquely distinguishes an individual in the pools.

**Figure 5:**
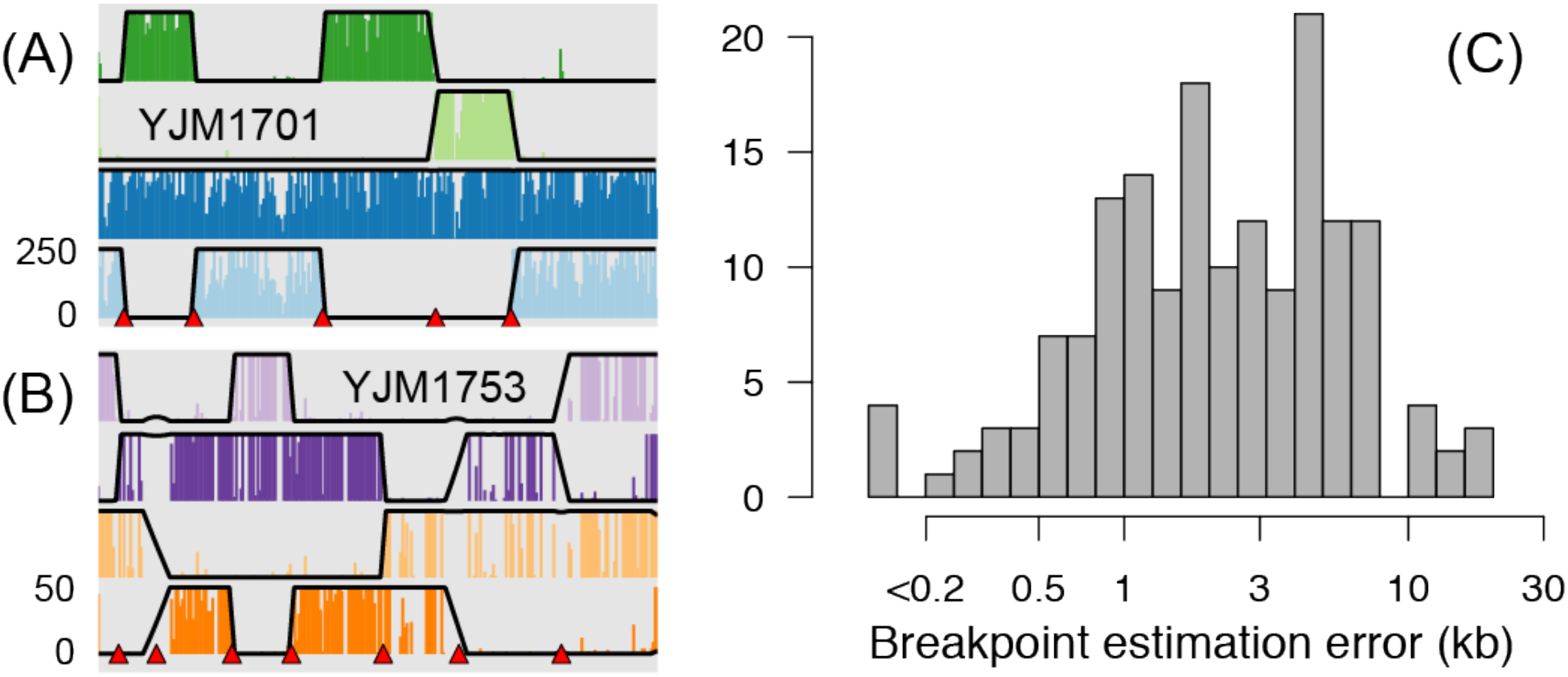
Read depth reveals blocks of ancestry inherited according to Mendelian expectations in the four spores resulting from a single meiosis. (A) Depth of uniquely mapping reads in 3kb bins tiling across chromosome II for four spores from a tetrad produced by a reference-YJM1701 hybrid. The four spores were each sequenced as members of different pools. Read depth plots across the chromosome are stacked for the four spores. Note the clear boundaries between blocks of ancestry inherited from parents and plainly visible breakpoints due to recombination. Read depths are trimmed at a maximum of 200 reads as indicated by axis labels on the left. Posterior probabilities of non-reference ancestry from the HMM are overlain on each read depth plot, scaled to the same range as the read depths. Red triangles indicate locations of inferred recombination breakpoints. (B) Read depth across chromosome II as shown in (A), but for a tetrad with a non-reference parent (YJM1753) that is among the least diverged in the pool, relative to the common reference. The read depth threshold is 20% that in (A) but segments of ancestry are still clearly visible and inferences of ancestry match intuition. (C) Histogram of the accuracy of recombination breakpoint estimation compared between pooled and unpooled data. This histogram is identical to that plotted in Figure 4B but using breakpoints inferred with the HMM that uses tetrad information.

The states in this model represent the six different ways in which parent *j* ancestry can be inherited in two of four segregants. The HMM directly models read counts in each bin such that, in each hidden state, we expect the bulk of reads to be divided equally between the two segregants in each cross that inherited that genomic segment from parent *j* and only a few reads (errors due to sequencing errors or mismapped reads) to map to the two segregants that inherited from the reference parent. We assume that crossovers are non-overlapping, constraining transitions between states to those that can be accessed by a single crossover. Figures 5A and 5B show ancestry probabilities derived from this HMM superimposed over the raw coverage data. It is apparent that ancestry calls match our intuition based on the coverage data.

As in our simpler model above, genotype probabilities across all segregants were strongly bimodal (Figure 5C), with >97% of genomic segments assigned a probability >99% of ancestry from one parent. Thus, genotypes can be confidently assigned as originating from one of two possible haplotypes for the vast majority of the genome. We compared recombination breakpoint calls using unpooled data as above, and found that incorporating meiotic expectations in the model further improved the accuracy and resolution of our calls. Specifically, our estimates using the pooled tetrad HMM were within 2.1kb (median) of the breakpoint locations called in unpooled data, as compared to 3.5kb (median) for the simpler HMM above. We also found fewer missed calls and false positive calls: of 169 breakpoints inferred using unpooled data, three were missed in pooled data and six incorrect breakpoints were called in the pooled data. Again, these breakpoints were either small blocks of ancestry (<15kb; *N=6)* or near (<40kb) the end of a chromosome (*N*=3). As above, these types of missed calls affected only a small portion of the complete genome (for these four segregants, 0%–0.6%).

### Cost savings

How cost-effective is the PHB approach for resolving genotypes? This question is highly dependent on the specific details of a particular experiment. For the yeast segregants detailed in this paper, we achieved ~92% savings in sample preparation costs by pooling using PHB (relative to non-pooled sequencing). This estimate excludes inhouse labor costs, which makes it highly conservative. Additionally, the PHB strategy greatly facilitates sample handling. In this example, we pooled cultures after growing cells to saturation, which allowed us to perform only 12 DNA extractions and 12 library preparations instead of the 96 that would be required for a strategy not utilizing pooling. Note that, while the example described above results in a small mapping population (*N*=96) that lacks substantial power for detecting and resolving QTL, the savings in library preparation costs and ease in sample handling scale linearly with size; thus, the same principles could be used to generate a far larger population.

### Conclusions

In this paper, we describe a method for pooled genotyping of offspring from multiple genetic crosses, such as those that that constitute multiparent populations. This method takes advantage of the fact that genotyping of multiparent populations can be conceptualized as tracing the inheritance of genomic segments of the population founders in recombinant offspring. We develop and test HMMs to genotype yeast segregants sequenced in a raw pool. These HMMs gave excellent accuracy and resolution for genotyping the F2 segregants we describe and may be useful for other applications as well. For some applications an even more powerful method would be to explicitly examine each private haplotype discriminating individuals in the pool and to utilize base quality scores and read mapping qualities to calibrate a probability that the haplotype was seen in the pool. Armed with this probability and an accurate genetic map, one could implement an HMM to infer genotype probabilities (Gatti *et al.* 2014) that avoids binning and counting reads altogether.

Although PHB could conceivably be applied in many different scenarios, it is clear that the strategy requires genetically diverse individuals in order to produce useful output. How diverse should the individuals composing each pool be? This question cannot be answered in general terms because the solution depends on biological factors specific to the organism and details of experimental design that are specific to the study. There are several factors important to any application of PHB, including the density of uniquely distinguishing polymorphisms across the genome, the expected haplotype block size, genome size, sequencing depth and read length, and the user’s desired confidence in genotype calls. Large haplotype blocks, a small genome, high read depth and long reads, and a high density (and relatively even distribution) of uniquely distinguishing polymorphisms will all result in improved ability to trace ancestry of genomic segments from parents to sequenced offspring. These issues are best resolved by conducting simulations of a short-read sequencing experiment using realistic sequencing error profiles and generating reads from offspring with a known genome derived via the desired breeding design, in order to empirically explore the accuracy of pooled genotyping using PHB. The strategy has built-in tunability in the sense that the pooling can be made more aggressive to maximize cost and labor savings or less aggressive to increase confidence in ancestry calls.

When does employing PHB make the most sense? The strategy is applicable for multiparent population designs where individuals are not exchangeable and can be pooled such that each individual in the pool has some unique ancestry. Given a compatible breeding design, the PHB strategy is most advantageous when library preparation constitutes a significant portion of the total project cost. This will be particularly likely for organisms with small- to medium-sized genomes, but will continue to be accentuated as sequencing throughput increases and costs fall. Furthermore, the strategy could be creatively applied to organisms with larger genomes in concert with techniques for reduced-representation genome sequencing (e.g.Peterson *et al.* 2012). PHB might be particularly pertinent for low budget scenarios such as investigators studying non-model systems or a single laboratory creating a mapping population to consider a question of particular interest. Aside from cost considerations, the PHB strategy also has potentially beneficial logistical advantages. In particular, for very large mapping populations (thousands of individuals) where sample handling is significant, PHB has the potential to reduce the number of samples by an order of magnitude or more.

## Methods

### Yeast crosses and generation of F2 segregants

We crossed a haploid, heterothallic ρ^0^ strain isogenic with S288c (hoΔ::kanMX4 MATα ρ^0^) to each of twelve other strains (hoΔ::loxP MATa ρ+; for details of strains used in this study, see Supplementary Table 1). We crossed a ρ^0^ strain to ρ^+^ strains to control for mitochondrial ancestry. In contrast to higher eukaryotes, offspring from a ρ^+^ × ρ^+^ S. *cerevisiae* cross possess highly recombinant mitochondria (Nunnari *et al.* 1997; Solieri 2010). A ρ^0^ × ρ^+^ cross thus allows for the control of mitochondrial ancestry and minimizes this potential confounding factor. All strains used (and thus the F2 strains generated in this study) lack auxotrophic mutations, which can be both debilitating and environmentally limiting.

We selected diploids and streaked for single colonies on YP(EG) (1% Yeast Extract, 2% Bacto Peptone, 1% ethanol, 1% glycerol, 2% Bacto Agar) + 200mg/L G418. In each tetrad we verified 2:2 segregation of G418 resistance by testing for growth on YPD (1% Yeast Extract, 2% Bacto Peptone, 2% dextrose, 2% Bacto Agar) + 200mg/L G418 and 2:2 segregation of mating type by PCR of the MAT locus. For each segregant, we also tested at least four restriction fragment length polymorphisms (RFLPs) that together uniquely distinguished pairs of the 13 strains studied, in order to verify expectations of ancestry and confirm 2:2 segregation of the RFLPs within each tetrad. The strain backgrounds that we chose for this study possess collinear genomes with no major translocations or known large inversions (Strope *et al.* 2015).

### Sample preparation for genotyping of yeast segregants

We revived segregants from frozen stocks and grew overnight in YPD at 30C. We transferred a toothpick of cells for each segregant to a different well containing 200ul YPD in a 96-well round-bottom plate. We grew these cultures for 48 hours to ensure saturation. To create pools we combined the full volumes of each well in every row of the plate, which resulted in eight pools. We extracted DNA from each pool using the Qiagen Genomic-tip kit. We performed library preparation using recommended Illumina protocols and sequenced 101 bp paired-end reads on an Illumina HiSeq machine.

### Read mapping, counting, and coordinate conversion

For all genomic analyses of yeast strains in this study, we used full genomes sequenced and *de novo* assembled by (Strope *et al.* 2015). We mapped short reads using bwa mem v0.7.12 (Li and Durbin 2009) and kept reads with a mapping quality of at least ten. We also tested mapping only perfectly matching reads using BBMap v35.x (https://sourceforge.net/projects/bbmap/) and found very similar overall results (data not shown). We sorted and manipulated bam files and marked duplicates using samtools v0.1.19 (Li *et al.* 2009) and Picard tools 1.101 (http://broadinstitute.github.io/picard). We used bedtools v2.25.0 to count reads mapping to bins of defined sizes (Quinlan and Hall 2010). To convert coordinates between the reference S288c genome and *de novo* assembled non-reference strain genomes we used liftOver (Hinrichs *et al.* 2006)

### Hidden Markov model for PHB genotyping

We implemented an HMM that provides probabilistic estimates of ancestry across each segregant’s genome that is sequenced in a pool. For a single segregant present in a pool, only reads that uniquely map to one of the *j* = 1,…,12 non-reference strains are visible evidence of inheritance from that segregant’s (known) parents, since reference strain ancestry is shared by all members of the pool (Figure 2). Thus, reference strain ancestry in a particular segregant must be inferred by the absence of reads uniquely attributable to parent *j* at a particular genomic locus. We examine read counts in 7.5kb bins tiling across the genome of each non-reference strain. Using the more sophisticated HMM incorporating tetrad information as described below, we examined a range of bin sizes and found the most stable estimates of recombination breakpoint count for bins between 5–10kb (Supplementary Figure 1).

Bins with many reads likely represent a genomic segment inherited from non-reference parent *j,* while bins with few reads likely represent either inheritance from the reference parent or bins with low mappability. This HMM has two underlying states – reference ancestry and parent *j* ancestry. In both states, reads are emitted according to the binomial distribution, with size equal to the total number of reads mapping uniquely to parent *j* across the genome and bin-specific "success” probabilities. In the reference ancestry state, reads are effectively invisible since this ancestry is not unique to the pool, so reads are emitted at bin-specific background error rates. In the parent *j* ancestry state, reads are also emitted at bin-specific rates. We used bin-specific probabilities because bins with low mappability are expected to have few uniquely mapping reads in either ancestral state, and factors such as sequence complexity and the density of private variation are expected to lead to large variability in read coverage density between bins. In order to calibrate these bin-specific probabilities as accurately as possible, we simulated reads from hybrid genomes using dwgsim v0.1.10 (https://github.com/nh13/DWGSIM) and obtained empirical estimates of the fraction of all uniquely mapping simulated reads within each bin. Specifically, we conducted multiple simulations where half of a hypothetical segregant’s genome was inherited from parent *j* and the other half from the reference strain, mapped reads to the pool genome using the same procedures as the real data, and tallied the fraction that mapped to each bin. For transitions between hidden states (switches in ancestry), we calculate a transition probability *c* (assumed to be constant across the genome) based on the size of genomic bins and an approximate average recombination rate in the yeast genome of one centiMorgan per two kilobases. For example, for windows of size 100bp, the transition probability between states separated by one recombination event would be 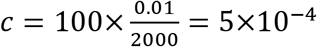.

We implemented the HMM described above using the python package hmmlearn v0.2.0 (https://github.com/hmmlearn/hmmlearn) and used this software to obtain posterior probabilities of ancestry for each bin. Before providing read coverage counts to the HMM we used the R package *mmand* (https://github.com/jonclayden/mmand) to apply a median filter of size 11 in order to sharpen edges at sites of breakpoints. After running the HMM and obtaining posterior probabilities of ancestry, we rounded probabilities to call blocks of ancestry and infer the locations of breakpoints. We filtered out any ancestry blocks supported by less than three bins as crossover interference suggests that such small blocks are likely spurious.

### Hidden Markov model for yeast tetrad genotypes

We implemented an HMM that utilizes information on segregant relatedness that is known due to tetrad structure. Since each tetrad results from a single meiosis, our expectation according to the principles of Mendelian inheritance is that parental alleles at any genomic locus should show 2:2 segregation in the offspring. This assumption is violated at sites of allelic gene conversion, but such conversion tracts should be relatively short and should not strongly influence our results.

As above, reads uniquely mapping to one of the *j* = 1, …, 12 non-reference strains are evidence of inheritance from that segregant’s non-reference parent, since reference strain ancestry is shared by all members of a pool, and reference strain ancestry must be inferred by the absence of reads uniquely attributable to parent *j* at a particular genomic locus. Our HMM is run separately for each of the 12 non-reference parents but simultaneously on data from four pools. Hidden states in the model represent the 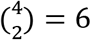 different possible ways in which the ancestry of two parents (A and B) can be partitioned 2:2 among the four spores in a tetrad: AABB, ABAB, ABBA, BAAB, BABA, and BBAA. Data emitted by each hidden state is four-dimensional and represents read counts (of uniquely mapping reads) across genomic bin *i* for each of the four spores in the tetrad. We model this data as multinomially distributed conditional on the total read coverage observed in bin *i.* Since we expect to observe only reads attributable to the non-reference parent, in the ideal case a bin with, say, 800 total reads would show ~400 reads in each of two pools and ~0 reads in the other two pools (data qualitatively matching this pattern is apparent in Figure 5A). In practice, mismapped reads and sequencing errors will result in some reads mapping to bin *i* even in the two pools without ancestry. Moreover, reads derived from pools with ancestry will not be split 50:50 due to differences in sequencing coverage between pools and variation in the abundance of each segregant within pools. Thus, we initialize emission probabilities randomly and optimize them as described below.

Transitions between states correspond to switches in patterns of ancestry that occur due to recombination events. For simplicity we assume zero probability of transitioning between states separated by more than one recombination event (e.g. AABB → BBAA). As above, for transitions between states separated by exactly one recombination event, we based the transition probability *c* on a rough estimate of the average rate of recombination in the yeast genome (1cM/2kb). From any of the six states, there is one inaccessible state (requiring two recombination events) and four accessible states (requiring one recombination event). Thus the probability of remaining in the current state is 1 – 4*c.*

Finally, we observed a chromosome-wide loss of heterozygosity in the two tetrads from the cross between YJM1617 and YJM1759, which we attribute to an event which occurred during the F1 mating of these two strains. Rather than discarding data arising from these segregants, we added a seventh hidden state where we expect to observe approximately equal read counts within bins tiling across an entire chromosome in offspring from this cross. We found that this method resulted in ancestry probabilities that matched intuition as well as results from segregants without loss of heterozygosity events (data not shown).

We implemented this HMM using hmmlearn v0.2.0 as above. For each tetrad, we ran the HMM at least six times using randomly initialized emission probabilities and used the Baum-Welch algorithm to optimize transition and emission probabilities. We used parameters from the model with the highest likelihood (typically, all runs produced likelihoods that were substantially similar) to calculate posterior probabilities of each hidden state for every genomic bin. Using this matrix of probabilities for each of the six states, the probability that each segment of a particular segregant’s genome is derived from parent *j* can be computed by summing posterior probabilities for all states matching this criterion. We filtered out any ancestry blocks supported by less than three bins as crossover interference suggests that such small blocks are likely spurious.

### Data and reagent availability

A pipeline to analyze the pooled sequencing experiment described in this study, as well as python code implementing the two HMMs described above, are available at github.com/daskelly/phb_paper. Yeast strains used in this study are available upon request.

## Acknowledgements

We gratefully acknowledge assistance from the North Carolina State University Genomic Sciences Laboratory, where library preparation and high-throughput sequencing were performed. We thank members of the Magwene lab, especially Debra Murray, for helpful comments during the conception and execution of this project.

